# Ancient BED-domain-containing immune receptor from wild barley confers widely effective resistance to leaf rust

**DOI:** 10.1101/2020.01.19.911446

**Authors:** Chunhong Chen, Bethany Clark, Matthew Martin, Oadi Matny, Brian J Steffenson, Jerome D Franckowiak, Martin Mascher, Davinder Singh, Dragan Perovic, Terese Richardson, Sambasivam Periyannan, Evans S Lagudah, Robert F Park, Peter M Dracatos

**Affiliations:** Agriculture & Food, Commonwealth Scientific and Industrial Research Organisation, GPO Box 1700, Canberra, ACT 2601, Australia; Plant Breeding Institute, The University of Sydney, Cobbitty, NSW 2570, Australia; Department of Plant Pathology, University of Minnesota, St. Paul, MN, USA 55108; Department of Agronomy and Plant Science, University of Minnesota, St. Paul, MN USA 55108; Leibniz Institute of Plant Genetics and Crop Plant Research (IPK) Gatersleben, 06466 Seeland, Germany; German Centre for Integrative Biodiversity Research (iDiv) Halle-Jena-Leipzig, Leipzig, Germany; Institute for Resistance Research and Stress Tolerance, Federal Research Centre for Cultivated Plants, Julius Kühn-Institute (JKI), Quedlinburg, Germany

## Abstract

Leaf rust, caused by *Puccinia hordei* is a devastating fungal disease affecting barley (*Hordeum vulgare* subsp. *vulgare*) production globally. Race-specific resistance (R) genes have been deployed widely; however, their durability is often compromised due to the rapid emergence of virulent *P. hordei* races, prompting the search for new sources of broad-spectrum resistance. Here we report on the cloning of *Rph15*, a broadly effective resistance gene derived from the wild progenitor *Hordeum vulgare* subsp. *spontaneum*. We demonstrate using introgression mapping, mutation and complementation that *Rph15* encodes a coiled-coil nucleotide-binding leucine-rich repeat (NLR) protein with an integrated Zinc-finger BED (ZF-BED) domain. The allelic variation at the *Rph15* locus was assessed using barley exome capture data that traced its origin to the western region of the Fertile Crescent bordering Jordan and Israel. To unravel the genetic relationship of two other leaf rust resistance genes (*Rph14* and *Rph16*) mapped at similar locus on chromosome 2H, we re-sequenced the *Rph15* gene from the near-isogenic line for *Rph15* (Bowman+*Rph15*) and the two donor accessions of *Rph14* (PI 584760) and *Rph16*, (PI 405292, Hs 680). Both whole genome and Sanger sequencing confirmed that Hs 680 carried *Rph15*, while *Rph14* in PI 584760 was an independent locus. A perfect diagnostic KASP marker was developed and validated to permit efficient introduction of *Rph15* into cultivated barley.

## Introduction

In 2016, barley (*Hordeum vulgare* subsp. *vulgare*) was ranked fourth among grain crops in production (141 million tonnes) behind maize, rice and wheat. It is used primarily for animal feed and malt production, but also serves as a major food staple in the mountainous areas of Central Asia, Southwest Asia, the Andes of South America and Northern Africa. Leaf rust, caused by the fungus *Puccinia hordei* Otth, is the most damaging and widespread rust disease of barley (Park et al. 2015). Leaf rust epidemics can occur globally and have been reported to cause significant reductions in grain quality and yield. Yield losses up to 62% have been reported in highly susceptible barley cultivars (Cotterill et al. 1992). Resistance to leaf rust in barley is controlled by qualitative type *<u>Rph</u>* (<u>R</u>eaction to *<u>P</u>uccinia <u>h</u>ordei*) genes conferring all-stage resistance (ASR) that is typically race-specific or by quantitative trait loci (QTL) conferring partial adult plant resistance (APR) that is generally not race-specific (Niks et al. 2015). Due to *P. hordei* evolution and the subsequent emergence of new pathogenic variants, ASR resistance to leaf rust is often transiently effective. Of the 22 catalogued ASR genes, five (*Rph10, Rph11, Rph13, Rph15* and *Rph16*) originate from *H. v*. ssp. *spontaneum* (Park et al. 2015).

The leaf rust resistance gene *Rph15* is derived from PI 355447, an accession of wild barley collected from Israel (Jin and Steffenson 1994; Chicaiza 1996). Remarkably, *Rph15* is broadly effective to a wide array (>350) of global *P. hordei* isolates, with virulence being found in only one isolate (90-3, from Israel) (B. J. Steffenson and T. G. Fetch, unpublished). *Rph15* therefore represents a valuable resistance source for disease control in cultivated barley especially when deployed in combination with other effective resistance genes. Chicaiza (1996) first determined that the *Rph15* resistance was inherited as a single dominant gene that mapped to the centromeric region of the short arm of chromosome 2H in close linkage to the RFLP marker MWG2133. Martin et al. (2020) recently developed a series of introgression lines (*Rph1-Rph15*) in the Bowman background and mapped *Rph15* to a physical interval (44-57Mb) on chromosome 2H in the Morex reference genome (Figure 1). Previous genetic studies also determined that two other widely effective *Rph* genes, *Rph14* and *Rph16*, were either allelic or very closely linked to *Rph15*, suggesting the presence of an allelic series or complex resistance locus on chromosome 2H (Ivandic et al. 1998; Weeraseena et al. 2004; Derevnina et al. 2014). Previous high-resolution mapping in the Bowman+*Rph15* x Bowman and Hs 680 x L94 mapping populations also determined that both *Rph15* and *Rph16* are either very closely linked or co-located with the RFLP marker MWG2133 (Perovic et al. 2004; Weeraseena et al. 2004) (Figure 1). In this study, our aim was to isolate *Rph15* and determine the molecular basis of the postulated allelic relationships among *Rph14, Rph15* and *Rph16* on chromosome 2H.

**Figure 1.**
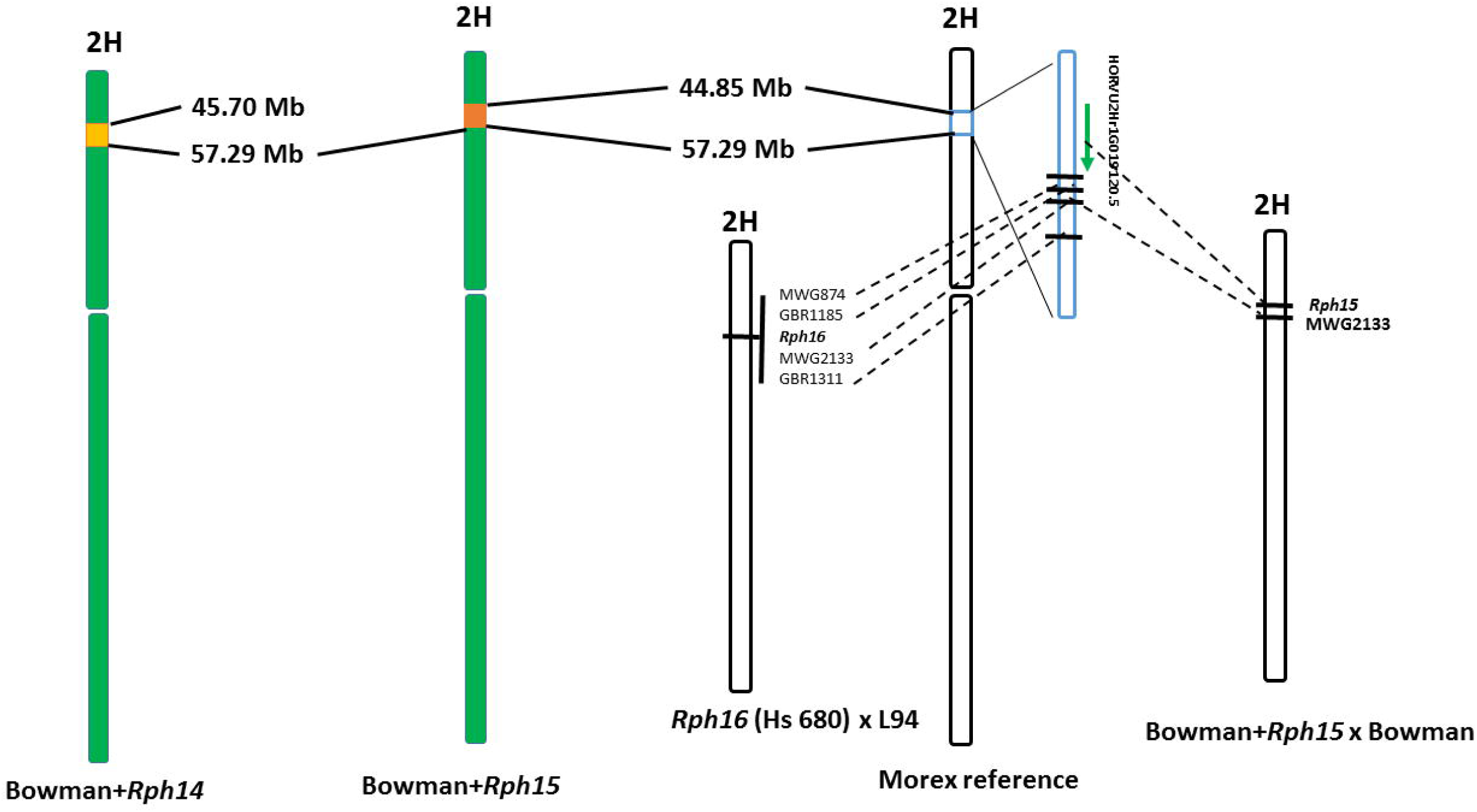
Comparative physical and genetic maps for chromosome 2H at the *Rph14/15/16* locus, including graphical genotypes for Bowman introgression lines carrying leaf rust resistance genes (A) *Rph14* and (B) *Rph15* from Martin et al. (2020) and the corresponding physical region in the Morex V1 reference genome assembly used to search for candidate NLR genes for further functional analysis. Different coloured sections within a chromosome are the regions of retained donor chromatin from PI 584760 (*Rph14.ab*) and PI 355447 (*Rph15*). Previously published genetic maps for chromosome 2H derived from mapping populations including: Bowman+*Rph15* x Bowman (Weeraseena et al. 2004) and HS084 x L94 (Perovic et al. 2004) used to map *Rph15* and *Rph16*, respectively, were included and co-segregating RFLP markers were anchored onto the Morex physical assembly using a Blastn algorithm in the IPK BLAST server.

## Results and Discussion

The previously reported monogenic inheritance and linkage of *Rph15* with RFLP marker MWG2133 was re-confirmed using a Bowman+*Rph15* x Gus F_2:3_ population (Supplementary File S1). To isolate the *Rph15* resistance gene, a candidate gene approach was adopted using the 2017 Morex reference assembly (Mascher et al. 2017) as a road map to search for immune receptors encoded by R genes, such as nucleotide binding site leucine-rich repeats (NLRs), within the target physical interval specified in Martin et al. (2020) (Figure 1). A single predicted NLR gene was identified (HORVU2Hr1G019120.5) and considered as the best candidate for further investigation using a loss of *Rph15* function mutant population generated by the treatment of Bowman+*Rph15* barley with sodium azide. Screening of progeny seeds from 4,320 M_2_ spikes with the *Rph15*-avirulent *P. hordei* race 5457 P+ identified eight independent mutants (*rph15*) which were confirmed at the M_3_ generation. A 9,442 base pair (bp) full-length genomic sequence of the candidate gene HORVU2Hr1G019120.5 that included 1.3 kb and 2.5 kb of 5’ and 3’ sequence, respectively, was amplified from *wt* Bowman+*Rph15* (Figure 2). We then sequenced the full-length gene from the eight mutants and identified non-synonymous mutations in six out of the eight lines including: a premature stop codon (M1695), serine to proline substitutions or *visa versa* (M1371-3, M1727, and M4022) and substitutions from glycine to either glutamic or aspartic acid (M4022 and M4321) (Figure 2). The remaining two susceptible mutants did not contain sequence changes in the candidate gene, suggesting they likely carry non-synonymous mutations within downstream signalling components essential for *Rph15*-mediated resistance.

**Figure 2.**
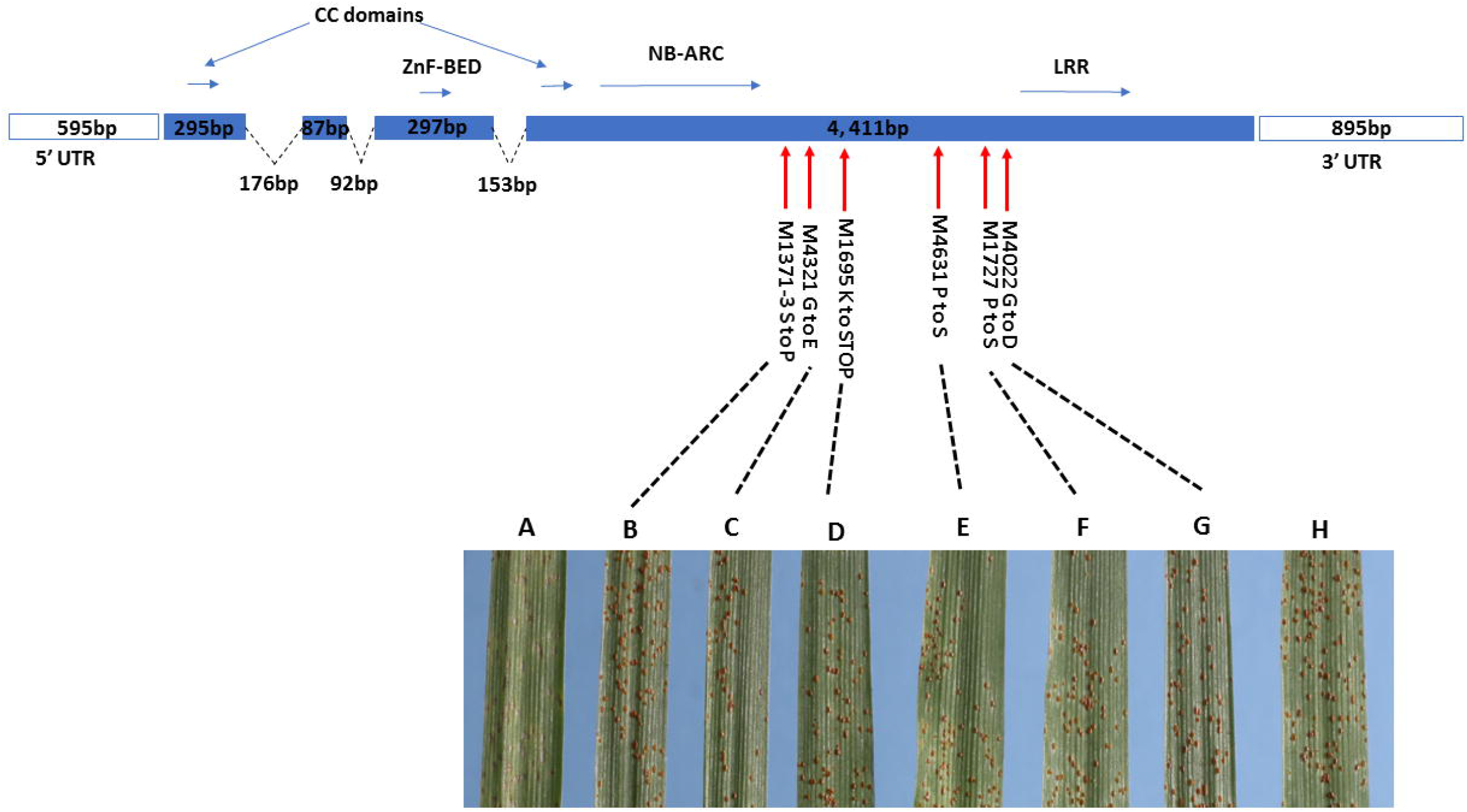
(A) Structure of the *Rph15* (HORVU2Hr1G019120.5) gene from the near-isogenic line Bowman+*Rph15*, including the coding region (solid blue) and 5′ and 3′ untranslated regions (UTRs, open blue boxes), and the position of the M1371-3, M1695, M1727, M4022, M4321 and M4651 mutations and the functional changes they induce resulting a loss of resistance. (B) Leaf rust infection phenotypes 12 days after inoculation of (L to R) wild type Bowman+*Rph15*, Bowman, mutants M1371-3, M1695, M1727, M4022, M4321 and M4651. For a full description of infection types refer to Park et al. (2015).

The full-length genomic sequence from Bowman+*Rph15* was cloned into the binary vector and transformed into the leaf rust susceptible barley cultivar Golden Promise. Six independent hemizygous T_0_ Golden Promise+*Rph15* transgenic lines were produced that contained the wild-type *Rph15* candidate in addition to two control lines that contained an empty vector only (Supplementary Figure S2). T_1_ progeny were evaluated at the seedling stage with Australian (5457 P+) and North American (92-7) *Rph15*-avirulent isolates and a rare *Rph15*-virulent *P. hordei* isolate (90-3) originally collected in Israel (Supplementary Figures S3 and S4). Four Golden Promise+*Rph15* T_1_ generation transgenic lines showed the expected segregation for low infection types (0; to ;1CN) characteristic of the *Rph15* resistance observed on Bowman+*Rph15* seedlings (Supplementary Figure S3). Two of the four transgenic lines showed complete susceptibility in response to the *Rph15*-virulent culture (90-3), whilst the segregation ratios in the remaining two T_1_ families unexpectedly segregated 1 : 3 <u>S</u>usceptible : <u>R</u>esistant. The observed resistance to the *Rph15*–virulent isolate may suggest auto-activation due to a gene dosage effect where over-expression of *Rph15* requires two or more copies of the transgene in the homozygous plants. To confirm the correlation between the presence of the transgene and the observed phenotypic response in the T_1_ transgenic families, 12 sib plants per T_1_ family were genotyped using a KASP marker designed to a SNP within the 3^rd^ intron of the *Rph15* candidate gene. Genotypic analysis confirmed, without exception, co-segregation between the presence of the heterozygous SNP genotype (Golden Promise susceptibility allele in combination with the T-DNA from Bowman+*Rph15*) in all resistant sibs, and the susceptibility allele found within all susceptible sib lines. Thus, the exploitation of previous mapping data, combined with both mutation and complementation experiments in the present study, facilitated confirmation of the candidate as the causal gene for *Rph15-*mediated resistance.

A recent study showed that four cloned wheat stem rust resistance genes (*viz. Sr22, Sr33, Sr35* and *Sr45*) transformed into barley conferred resistance to *P. graminis* f. sp. *tritici* (*Pgt*; causal agent of stem rust), but not to the barley-adapted leaf rust pathogen *P. hordei* (Hatta et al. 2018). We transformed the rust susceptible wheat cv. Fielder with the *Rph15* gene driven by its native promoter and tested its effectiveness against the wheat-adapted leaf rust (*P. triticina*) and stem rust (*Pgt*) pathogens. All wheat Fielder+*Rph15* transgenics were as susceptible to both rust pathogens as the Fielder control (data not shown). These data indicate that either *Rph15* is non-functional in wheat or alternatively *P. triticina* and *Pgt* do not produce a corresponding effector recognised by *Rph15*.

Among myriad of molecular functions hidden in plant NLRs, the kinase, WRKY and BED domains are most frequently detected (Kroj et al. 2016). Rare sub-families of NLR immune receptors in plants have evolved to encode additional integrated domains that act as decoys to recognise and interact with pathogen effectors that would normally target host transcription factors containing Zinc Finger-BED (ZF-BED) domains (Grund et al. 2019). The ZF-BED domain, originally characterised in Drosophila through mutagenesis studies, was shown to be essential in rice and wheat to confer *Xa1* and *Yr7*-mediated resistance, respectively (Yoshimura et al. 1998; Marchal et al. 2018). Polymerase chain reaction (PCR) using cDNA indicated that the *Rph15* gene consists of four (three small and one large) exons and encodes an immune receptor with a single predicted BED-domain nestled between two coiled-coil domains followed by the canonical NLR (1696 residues) (Figure 2). Three BED domain NLRs (*Yr5, Yr7* and *YrSP*) with unique pathogen specificity conferred resistance to stripe rust in bread wheat on the long arm of chromosome 2B, comprising a complex resistance NLR cluster (Marchal et al. 2018). The presence of four exons and only a single BED domain contradicts the characteristic three exon gene structure commonly identified for BED domain-containing NLRs within the syntenic region of the *Yr5/Yr7/YrSP* locus recently characterised in bread wheat (Marchal et al. 2018). Phylogenetic analysis using numerous recently cloned NLR immune receptors from the Triticeae not surprisingly determined *Rph15* was most closely related to other BED-domain carrying NLR proteins including both *Yr5* and *Yr7* homologues from wheat and *Xa1* from rice on chromosome 4 (corresponding to 2H in barley) (Supplementary Figure S5). Interestingly, *Rph15* clustered with previously cloned NLRs from bread wheat conferring both rust and mildew resistance relative to the MLA clade and the recently cloned leaf rust resistance NLR *Rph1* from cultivated barley (Dracatos et al. 2019; Supplementary Figure S5).

According to Marchal et al. (2018), *Yr5* and *Yr7* carried BED domains that clustered with other clade I BED domain carrying NLRs from the Triticeae, whilst *Xa1* from rice clustered separately from both clades I and II yet still carried the nine conserved residues between BED I and II clades. Further examination of the BED domains between *Rph15, Yr5/Yr7* and *Xa1* suggest that the BED domain from *Rph15* was more closely related to the BED II clade consensus and yet only carried six out of the nine conserved residues, suggesting the possible presence of functionally diverse BED domain of distinct from BED I and II. Although no mutants were identified within the BED domain of *Rph15* in this study, our yeast-two-hybrid data confirmed the strongest transcriptional activation using the CC-BED-CC construct, whilst weak activation was also observed in the BED-CC relative to other *Rph15* protein domain configurations (Figure 3). Interestingly, no activation was observed using the BED domain of *Rph15* alone, suggesting that coiled-coil domains are also required for activation (Figure 3). The retained functional capacity of the ZF-BED domain in the *Rph15* protein supports its origin from a transcription factor and the evolution of the decoy model in plant NLRs. The alternative hypothesis is that there is no direct interaction of the ZF-BED and the pathogen effector. Rather, the transcriptional function we observed for the ZF-BED domain translates to its interaction with a partnering protein within the complex, providing the necessary transcription machinery for pathogen recognition.

**Figure 3.**
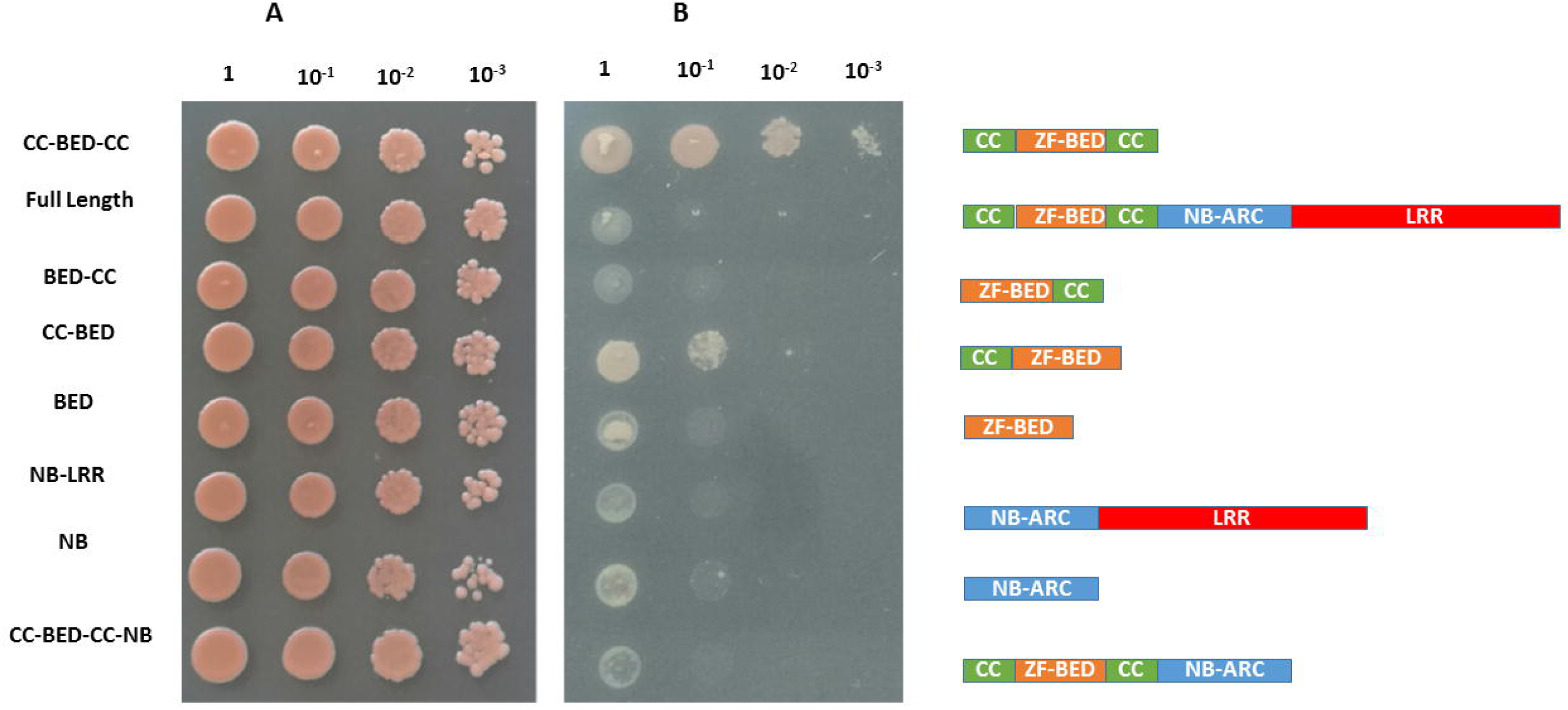
Yeast-two hybrid assay to test the transcription activation of a selectable marker to permit growth on selection media lacking Histidine using different domain configurations from within the RPH15 protein including: CC-ZF-BED-CC, ZF-BED-CC, CC-ZF-BED, ZF-BED, full length RPH15 protein, NBS-LRR, NBS, and CC-ZF-BED-CC-NBS. The different domain constructs were transformed into yeast and inoculated using 10-fold series dilutions on selection media either lacking either (A) Tryptophan (SC-T) or (B) Tryptophan and Histidine (SC-TH) and cultured at 30°C for 3 days.

The level of diversity associated with the *Rph15* allele is similar to that observed with barley genes that originated before the domestication of barley. Martin et al. (2020) reported on the development and characterisation of numerous *Rph* introgression lines in the Bowman background (BW lines). Three BW lines with the *Rph15* allele have a 2HS / 4HS reciprocal translation with the breakpoint in 2HS distal to the *Rph15* locus. The three BW lines (BW723 from PI 405179; BW724 from PI 405341; and BW725 from PI 391004) have two different marker patterns distal from the *Rph15* locus. These BW lines have identical marker haplotypes in the interstitial region between the breakpoint and the centromere where viable recombinants are extremely rare. The *Rph15* locus is in this interstitial region of 2HS. Because the *Rph15* resistance allele in the 2HS / 4HS translocation cannot be reassociated with different molecular markers by recombination, the *Rph15* mutant arose before the translocation. These observations suggest that the *Rph15* resistant mutant is very old, likely before the domestication of barley. There is, however, no evidence that the critical allele (*Rph15*) for *P. hordei* resistance at the *Rph15* / *Rph16* loci originated more than once. To determine the likely origin of the *Rph15* resistance, we assessed allelic diversity using exome capture data from a previously characterised panel of 267 accessions comprised of both wild and cultivated barley genotypes (Russell et al. 2016). None of the wild accessions carried the *Rph15* allele. However, based on sequence similarity of the five closest neighbouring accessions (80-90% nucleotide similarity), we postulate the origin of the resistance haplotype to be in the Western Fertile Crescent bordering Israel and Jordan (Figure 4).

**Figure 4.**
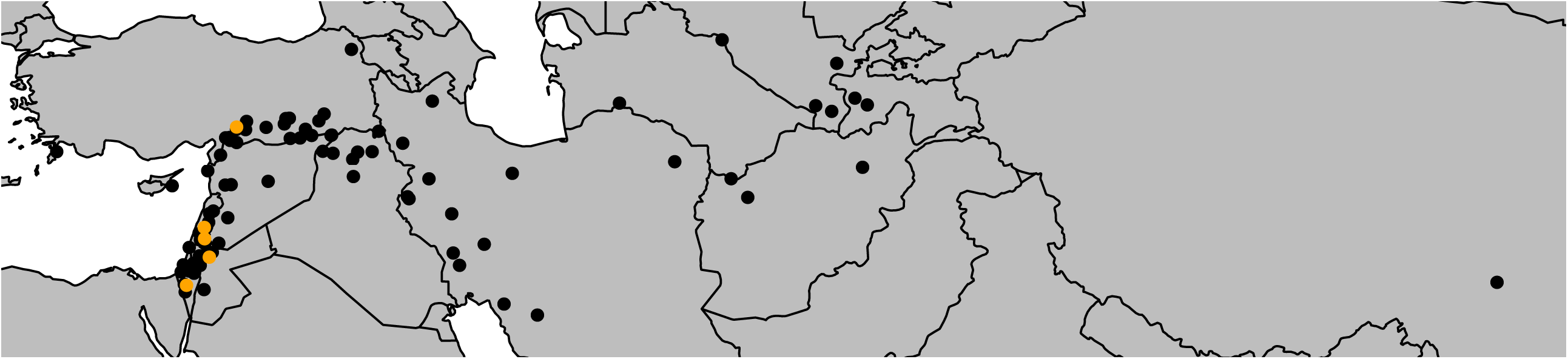
The geographic distribution of 91 wild barleys (*Hordeum vulgare* ssp. *spontaneum*) from the exome capture based on collection sites from the Fertile Crescent as given by Russell et al. (2016). The five nearest neighbours to the *Rph15* genomic sequence are shown by orange dots, while the other more distantly related haplotypes are denoted by black dots.

The presence of NLR clusters and multiple disease resistance alleles with different pathogen specificities arising through duplication and co-evolution is well established in the plant kingdom (Michelmore and Meyers 1998; Dracatos et al. 2019). Three leaf rust resistance loci, (i.e. *Rph14, Rph15* and *Rph16*) assumed to be independent have been mapped to the same chromosomal region on the short arm of 2H. Previous genetic studies suggest that *Rph15* and *Rph16* are either very tightly linked, alleles of the same gene or possibly conditioned by the same resistance allele. In contrast, according to Derevnina et al. (2014), *Rph14* and *Rph15* appear to be closely linked independent genes. To further unravel the genetic relationships among *Rph14, Rph15* and *Rph16*, we performed Illumina whole genome sequencing (WGS) in all three resistance donors, PI 584760, Bowman+*Rph15* and Hs 680. We then replaced the *rph15* susceptibility allele in Morex with the *Rph15* resistance allele sequence and mapped the Illumina reads of the sequenced accessions and then called the SNP variants. Despite incomplete coverage based on high sequence divergence, we determined that the *Rph14* resistance is due to a gene independent of *Rph15.* Sequence comparison of the *Rph15* gene from both Bowman+*Rph15* and the putative *Rph16* donor accession Hs 680 using both Illumina and Sanger methods revealed no nucleotide polymorphisms across the entire *Rph15* coding sequence. Further examination of historical phenotypic data using six North American *P. hordei* races determined that *Rph15* and *Rph16* shared the same race specificity; however, *Rph14* exhibited a different specificity in comparison (Supplementary Table 1). We therefore propose that 1) *Rph14* is independent from *Rph15* and 2) wild barley accessions (named Hs 680 and HS084) previously thought to carry the independent resistance locus *Rph16* in fact carry the same resistance allele as Bowman+*Rph15*.

Given that the number of disease resistance genes in plant species is finite, it is crucial to use these genes with appropriate stewardship to ensure they remain effective for as long as possible. Experience has shown that the most effective strategy to protect resistance genes and achieve durable disease resistance in crop species such as wheat (Park 2007) and barley (Park et al. 2015) is to deploy multiple genes in combination. Effective gene deployment however is reliant on the presence of diagnostic molecular tools to reduce the hitchhiking effect of deleterious traits or alternatively pyramiding resistance genes into cassettes using a transgenic approach. We developed a KASP marker by interrogating a synonymous SNP within the 3^rd^ intron of the *Rph15* gene and assessed its utility across 61 resistant near isogenic lines in the Bowman background (BW lines) (Martin et al. 2020). Based on data reported in Martin et al. (2020), from the 61 BW lines tested for rust resistance, 50 were postulated to carry *Rph15* either singly or in combination with another resistance gene and the remaining 11 carried unknown resistance (Supplementary Table 2). Interestingly, six of the BW lines postulated to carry *Rph15* and a further six BW lines with unknown leaf rust resistance carried a different SNP marker haplotype to either Bowman or Bowman+*Rph15*, suggesting they either may carry uncharacterised resistance or allelic variants of the *Rph15* gene (Supplementary Table 2). Eighty Australian cultivars with contrasting leaf rust response were also genotyped with the KASP marker, and as expected based on seedling leaf rust tests, all lacked, the *Rph15* marker haplotype was absent.

Since *Rph15* is effective against a global collection of over 350 *P. hordei* isolates (B. J. Steffenson and T. G. Fetch, unpublished) and is now tagged with a perfect KASP marker, it will be valuable for the efficient development of resistant barley cultivars through molecular breeding. It is possible that the widespread effectiveness of *Rph15* is due to its limited deployment in cultivated barley. To date, *Rph1*5 has been deployed in only one barley cultivar (ND Genesis) (J. D. Franckowiak and B. J. Steffenson, unpublished). Nevertheless, some NLRs have remained durable when deployed singly, especially when deployed in combination with one or more minor resistance genes. As more resistance genes and their cognate avirulence genes are cloned and resistance mechanisms and protein structures determined, more signatures of durability are likely to become evident. Further work is underway to determine the target of the ZF-BED domain to unravel the underlying mechanism of the *Rph15* resistance.

## Methods

### Plant materials and pathogen isolates

We used the near-isogenic line for *Rph15*, Bowman+*Rph15 (*AUS 490745; PI 355447/7*Bowman), originally sourced from Brian Steffenson as the resistance donor wild type for the development of mutants, population development and DNA template for complementation experiments. A low resolution F_2_-F_3_ mapping population was developed by crossing Bowman+*Rph15* with the leaf rust susceptible cv. Gus to re-confirm the chromosome arm location and the inheritance of the *Rph15* resistance. Complementation analysis in barley and wheat was performed using cvs. Golden Promise and Fielder, respectively. A detailed list of the rust pathogen isolates used in this study and their virulence phenotypes can be found in Supplementary Table 3.

### Candidate gene identification and amplification from Bowman+*Rph15*

Based on the hypersensitive infection type of the *Rph15* resistance, we used the physical interval of 44-57Mb reported in Martin et al. (2020) for the introgression of the *Rph15* resistance from PI 355447 into Bowman to search for predicted annotated NLR candidate resistance genes in the Morex reference assembly https://apex.ipk-gatersleben.de/apex/f?p=284:10. We designed a single primer pair (RGA4 F: and RGA4 R) based on the Morex *rph15* pseudogene sequence (GenBank: AY641411.1) to amplify a 9,442 base pair genomic fragment including approximately: 1.3 kb of sequence upstream from the translation Start codon and 2.5 kb of sequence downstream of the STOP codon. Polymerase chain reaction was performed using Phusion polymerase as per the manufacturer’s instructions (New England Biosciences). The PCR products were separated on a 1% agarose gel and a single band of the expected size was excised and purified. The gel-purified PCR product was then cloned into the sub-cloning vector TOPO XL using manufacturer’s instructions (Invitrogen). We amplified and cloned genomic fragments from resistant (Bowman+*Rph15* and Hs 680) and susceptible (Gus and L94) barley accessions. The plasmid DNA of five positive clones from each amplicon was sent for Sanger sequencing using internal primers and the sequences were compared with the template from Morex.

### cDNA analysis

Total RNA was extracted using the TriZol method (Invitrogen) as per manufacturer’s instructions. cDNA synthesis, 5’ and 3’ RACE (rapid amplification of cDNA ends) were performed using the manufacturer’s instructions (Clontech).

### Generation of loss of function mutants for *Rph15*

We performed sodium azide mutagenesis on the wild type *Rph15* donor barley line Bowman+*Rph15* (AUS 490745; PI 355447/7*Bowman) using the procedure described by Chandler and Harding (2013) with some modifications. Approximately 1,500 seeds were immersed in water at 4°C overnight. The imbibed seeds were transferred to a 2-litre measuring cylinder filled with water and aerated with pressurised air for 8 h, with one change of water after 4 h. The water was drained and the seeds were incubated in a shaker for 2 h in freshly prepared 1 mM sodium azide dissolved in 0.1 M sodium citrate buffer (pH 3.0). Next, the seed was washed extensively in running water for at least 2 h, and placed in a fume hood to dry overnight. Seeds were sown in the field and single spikes from each plant were harvested separately from the remaining spikes, which were harvested in bulk.

The Bowman+*Rph15* mutant M_2_ spikes and M_2_-derived M_3_ families were phenotypically assessed at the seedling stage as described by Dracatos et al. (2019). In all cases at least two susceptible plants were transplanted for each candidate M_2_ family segregating for *rph15* (*Rph15* knockouts) for progeny testing. Sequence confirmation for each mutant was performed through PCR amplification of M_3_ derived susceptible progeny for each mutant family.

### Sequence confirmation of *rph15* mutants

Sanger sequence confirmation of all mutants was performed at the M_3_ stage using a three-step process. Firstly, only DNA from progeny-tested homozygous susceptible families was extracted using the CTAB method (Doyle and Doyle 1987) for PCR amplification of the 9,442 bp genomic fragment for the candidate *Rph15* gene as described above. Secondly, the PCR products were cloned into the TOPO vector as described above and three positive clones for each amplicon were sent for Sanger sequencing for comparison with the wild type *Rph15* candidate gene. Finally, only when all three clones carried the same non-synonymous sodium-azide induced (either G to A or C to T) mutation were they deemed confirmed mutants.

### Diagnostic SNP marker development for *Rph15* and validation

A diagnostic Kompetitive Allele Specific PCR (KASP) marker assay was designed by interrogating a C/G SNP within the 2^nd^ intron of the *Rph15* gene. KASP assay was set up in 8 µl, including 4 µl of KASP master mix (LGC), KASP Assay mix (0.12 pmol two allele-specific primers and 0.3 pmol common reverse primer) and 25 to 50 ng of genomic DNA. KASP was conducted as follows: 1 cycle at 94°C for 15 min, 10 cycles of at 94°C for 20 sec and 65°C for 1 min, and then 32 cycles of at 94°C for 20 sec and 57°C for 1 min, cooled down to 25°C for 5 min, before reading the plate. Marker validation was performed using a panel of 80 Australian barley accessions all postulated to lack *Rph15* and 60 leaf rust resistant Bowman near-isogenic lines of which many were postulated to carry the *Rph15* resistance. All accessions are listed in Supplementary Table 3.

### Agrobacterium mediated transformation (ABMT)

#### Barley

The 9,442 bp fragment containing its native promoter and terminator was amplified by PCR with *NotI* at both sites, enzyme cutting and ligation into pWBVec8^15^. Subsequently, the vector containing intact sequence-verified *Rph15* candidate gene was transformed into *Agrobacterium tumefaciens* strain AgL0 using the freeze/thaw method (Chen et al. 1994). Barley transformation into Golden Promise was performed as described by Harwood et al. (1994).

#### Wheat

The *Rph15* gene fragment cloned into the binary vector pWBVec8 was transformed into *A. tumefaciens* strain GV3101. ABMT in wheat cv. Fielder was performed as described in Richardson et al. (2014). T_0_ plants were assessed for response to *P. graminis* f. sp. *tritici* race 98-1,2,3,5,6, *P. triticina* race 10-1,3,9,10,11,12, and *P. striiformis* f. sp. *tritici* race 134 E16 A+ at the 2^nd^-3^rd^ leaf stage as described by McIntosh et al. (1995).

#### Nearest neighbour analysis using the *Hordeum* exome capture

The genomic sequence of *rph15* was aligned to the genome sequence assembly of barley cultivar Morex (Morex V1, Mascher et al. 2017) with BWA-MEM (Li 2013) using default parameters. Sequence variants between Morex and *rph15* were detected with BCFtools mpileup (Li 2011). The results were imported into R (R Core Team 2016) and intersected with a SNP matrix derived from exome capture data of 267 wild and domesticated barley accessions (Russell et al. 2016). Nearest neighbors of *rph15* in a set of 91 wild barleys were found by a simple SNP matching distance. A map showing the collection sites of the 55 neighbors was plotted using the R package mapdata (https://cran.r-project.org/web/packages/mapdata/index.html). Collection sites were taken from Russell et al. (2016).

### Yeast two hybridisation

#### Cloning

The fragments of *Rph15* full length, *Rph15*-CC domain (Rph15-CC-BED-CC), *Rph15*-BED-CC, *Rph15*-CC-BED, *Rph15*-BED, *Rph15*-NBS-LRR, *Rph15*-NBS, and *Rph15*-CC-BED-CC-NBS (Primer information is detailed in Supplementary Table 4) were amplified by PCR using Phusion polymerase as per the manufacturer’s instruction. The fragments were separated using a 1% agarose gel, excised and cloned into pDON207 by Gateway BP reaction (Invitrogen). LR reactions were conducted to clone the different PCR fragments representing the domain configurations described above. Cloning was performed using the pDON207 vector and pGBKpGBK-NSC to generate plasmids BD-*Rph15*-Full length, BD-*Rph15*-CC-BED-CC, BD-*Rph15*-CC-BED, BD-*Rph15*-BED-CC, BD-*Rph15*-BED, BD-*Rph15*-NB-LRR, BD-*Rph15*-NB and BD-*Rph15*-CC-BED-CC-NB. DNA from each plasmid was sent for Sanger sequence confirmation prior to transformation into yeast.

### Yeast transformation

The transformation of each plasmid for the respective domain configurations was performed as described by Gietz and Schiestl (2007) using *Saccharomyces cerevisiae* strain HF7C (Clontech).

### Transcription activity assay

The yeast-two hybrid assay was based on the Gal4 system. Because the plasmid pGBK-NSC contains a tryptophan-producing gene, the transformants can be selected on the medium without tryptophan (SC-T). All the plasmid DNA was transformed into yeast and grown at 30°C on SC-T. If the peptide fusing with bind domain (BD) possesses transcription activity, the fusing protein can bind upstream of the histidine gene in yeast nuclei and activate the transcription of histidine, resulting in yeast growing on the selection medium without histidine (SC-H). Therefore, a single colony on the SC-T medium was selected and resuspended in sterilised water. A series of 10-fold dilutions were made and cultured at 30°C on SC-T and SC-TH (selection medium without tryptophan and histidine) in parallel. An image was taken after culturing for 3 days.

### Illumina Whole Genome Sequencing

A total of 5 µg of genomic DNA from each of the *Hordeum* stock accessions for genes *Rph14* (PI 584760), *Rph15* (Bowman+*Rph15*) and *Rph16* (PI 405292-Hs 680) was sent to Novogene Illumina (Hong Kong) for whole genome sequencing on a fee for service basis. We sequenced both Bowman+*Rph15* and Hs 680 80Gb of data (corresponding to roughly) 40x coverage using 2 x 250PE reads on the NovaSeq based on the proprietary protocols specified in the Illumina website. For *Rph14* (PI 584760), we sequenced using 2 x 150PE reads at 10 x coverage. Reads were mapped with Minimap2 to a modified Morex reference genome in which the closest homolog of *Rph15* had been replaced with the *Rph15.ad* allele sequence amplified from Bowman*+Rph15* (Li 2018). Variant calling and read depth computation were done with SAMtools.

### Phylogenetic analysis

The predicted *Rph15* amino acid sequence (HsRph15) was used as a query in GenBank using the program BlastP to identify closely related sequences. The HsRph15 amino acid sequence was then compared with that of related NLR sequences from *Oryza sativa* (Os) Xa1 (BAM17617.1) Triticeae including: *Aegilops tauschii* (Ata) Sr33-AGQ17384.1), *Secale cerale* (Sc) Sr50-ALO61074.1), *Triticum aestivum* (TaYr5, TaYr7, Ta;Lr22a-ARO38244.1, Sr22-CUM44212.1, Sr45-CUM44213.1, Pm2-CZT14023.1 and Pm8-AGY30894.1), (Tm; Sr35-AGP75918.1 and MLA1-ADX06722.1), *T. dicoccoides* (Td; Lr10-ADM65840.1) and *Hordeum vulgare* (Hv; Rph1-MK376319, MLA1-AAG37354.1, MLA6-CAC29242.1, and MLA9-ACZ65487.1). An unrelated NLR from Arabidopsis (*Arabidopsis thaliana*), At5g45510-Q8VZC7.2, was included as an out-group. A multiple sequence alignment was performed using ClustalW (Larkin et al., 2007), in Geneious version 11.0.2 (https://www.geneious.com) with the BLOSUM scoring matrix and settings of gap creation at 210 cost, and gap extension at 20.1 cost per element. After removing all ambiguously aligned regions using trimAl (Capella-Gutiérrez et al., 2009), the final sequence alignment of length 1,826 amino acids (n= 18) was determined. A phylogenetic tree based on this alignment was then inferred using the Neighbor-Joining method in the Geneious Tree Builder software, employing the Jukes-Cantor genetic distance model. Bootstrap support for individual nodes was generated using 1,000 bootstrap replicates.

## Acknowledgements

This work was supported by funds provided through Grains and Development Corporation (US00074), Judith and David Coffey and family, and the Lieberman-Okinow Endowment at the University of Minnesota. We are grateful to Matthew Williams (University of Sydney), Smitha Louis and Dhara Bhatt (CSIRO Agriculture and Food, Australia) for technical assistance. The authors also thank Dr Peter Dodds for commenting on the manuscript.

## Author contribution

C.C., P.M.D., B.C., O.M., Martin, M., Mascher, M., T.R., J.F., B.J.S., S.P., performed experiments. P.M.D., D.S., D.P., R.F.P., M.A., J.F., E.S.L., B.J.S., provided essential biological material for the study and assisted in the interpretation of the data. R.F.P, and E.S.L, provided infrastructure and funding for the project. P.M.D., wrote the manuscript with contribution from C.C. All authors approved the final manuscript before submission.

