## Supplementary figures and images for "Ancient BED-domain-containing immune receptor from wild barley confers widely effective resistance to leaf rust"

### Supplementary Figure S1

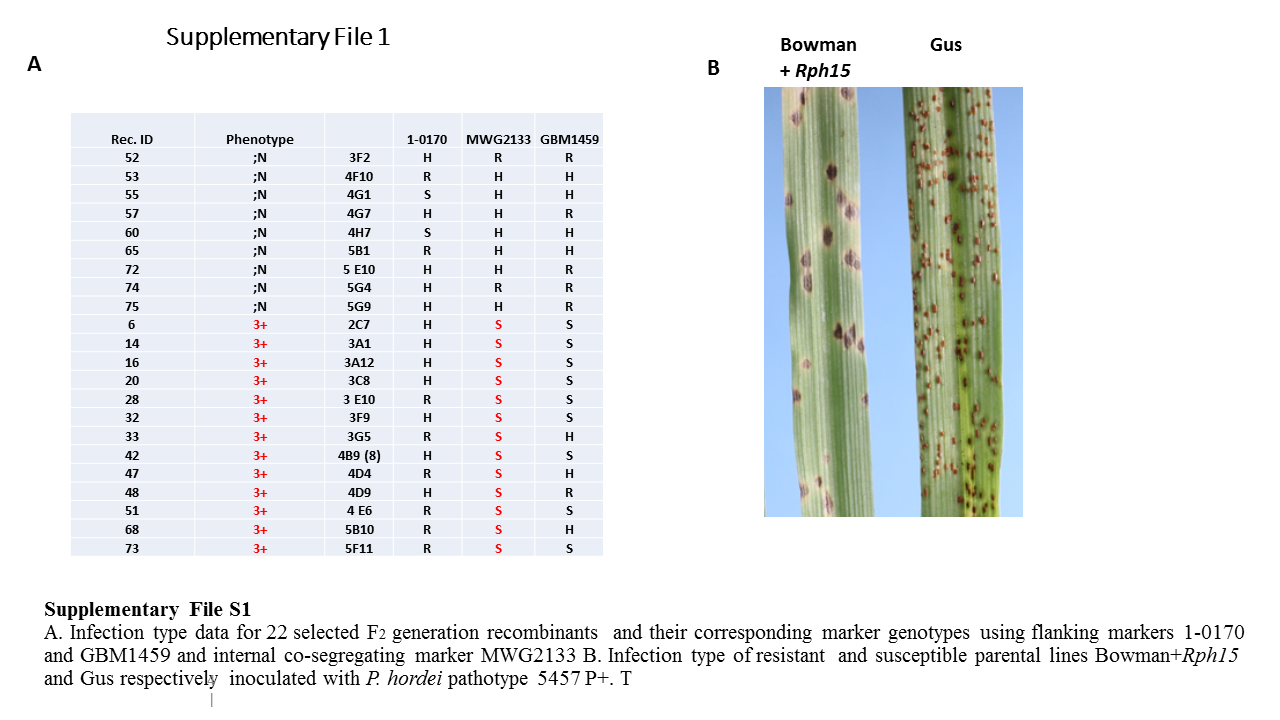

### Supplementary Figure S2

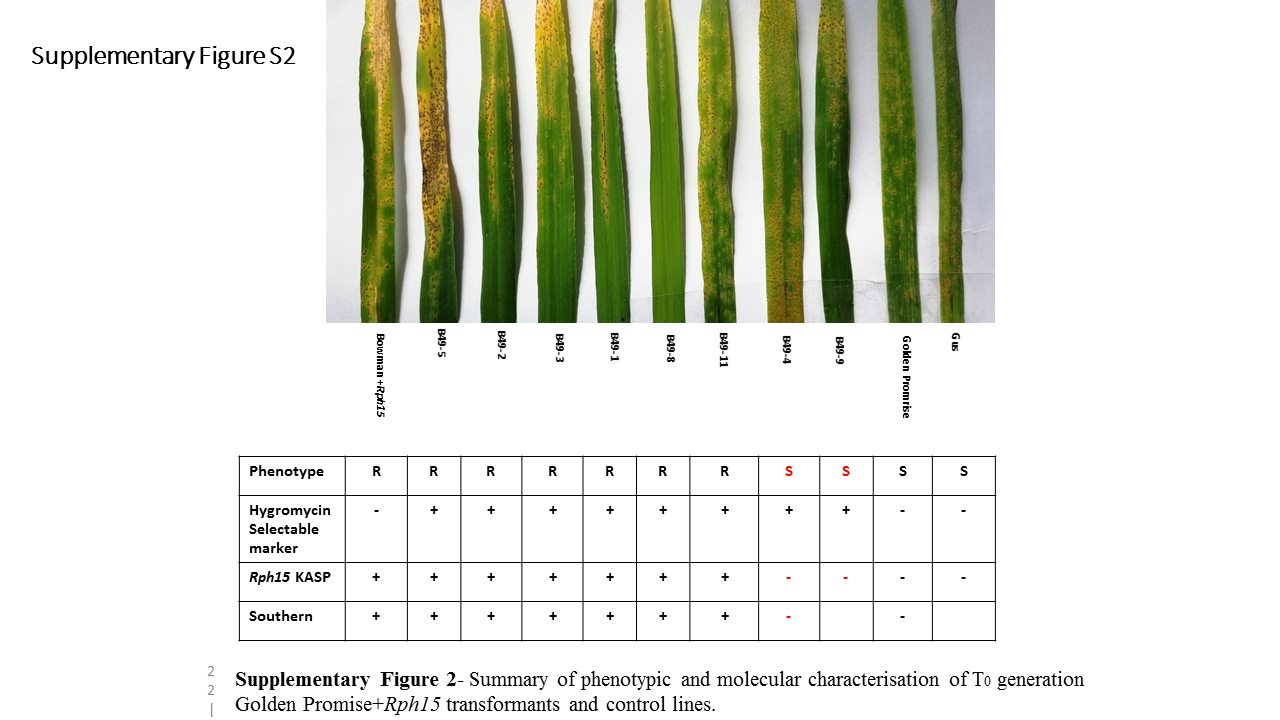

### Supplementary File S3

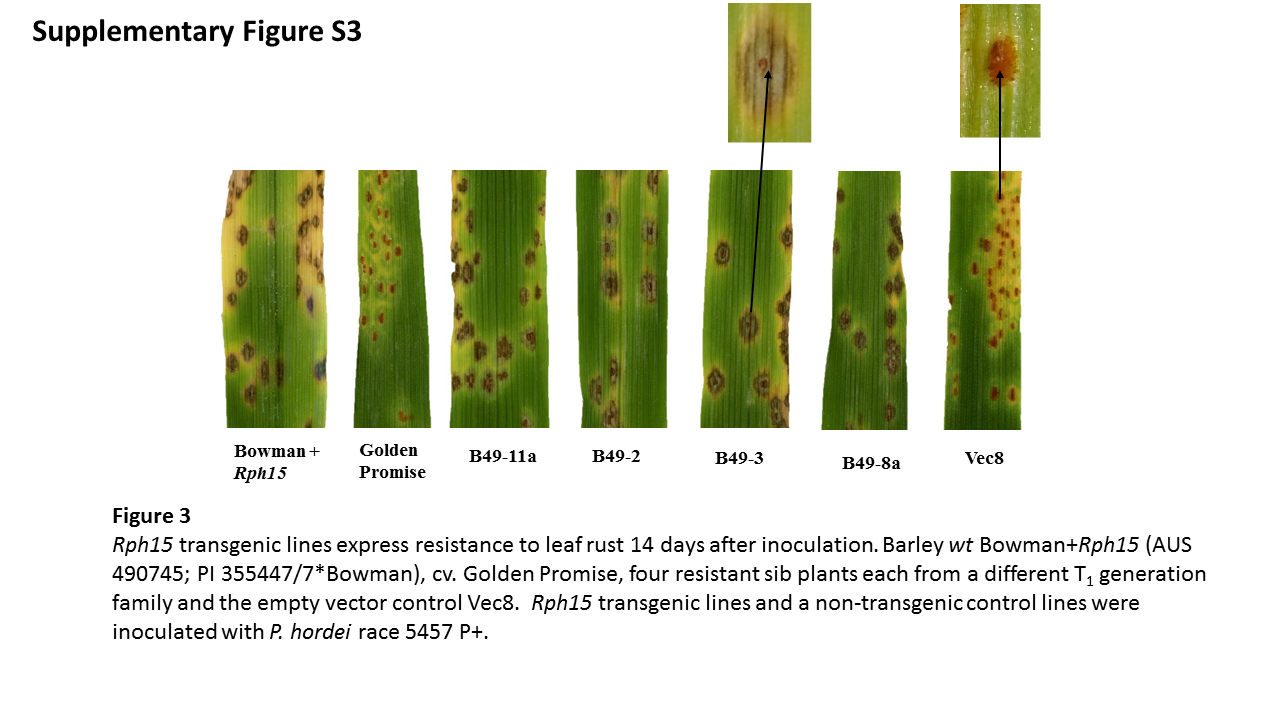

### Supplementary File S4

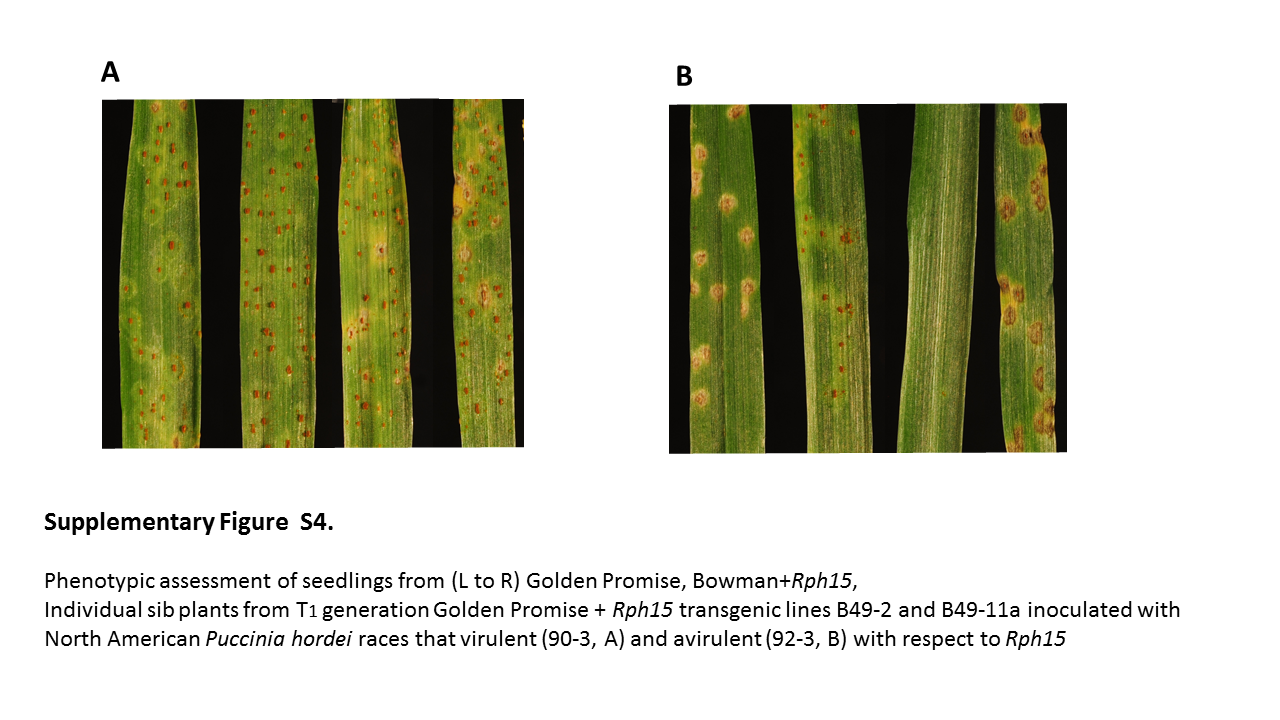

### Supplementary File S5

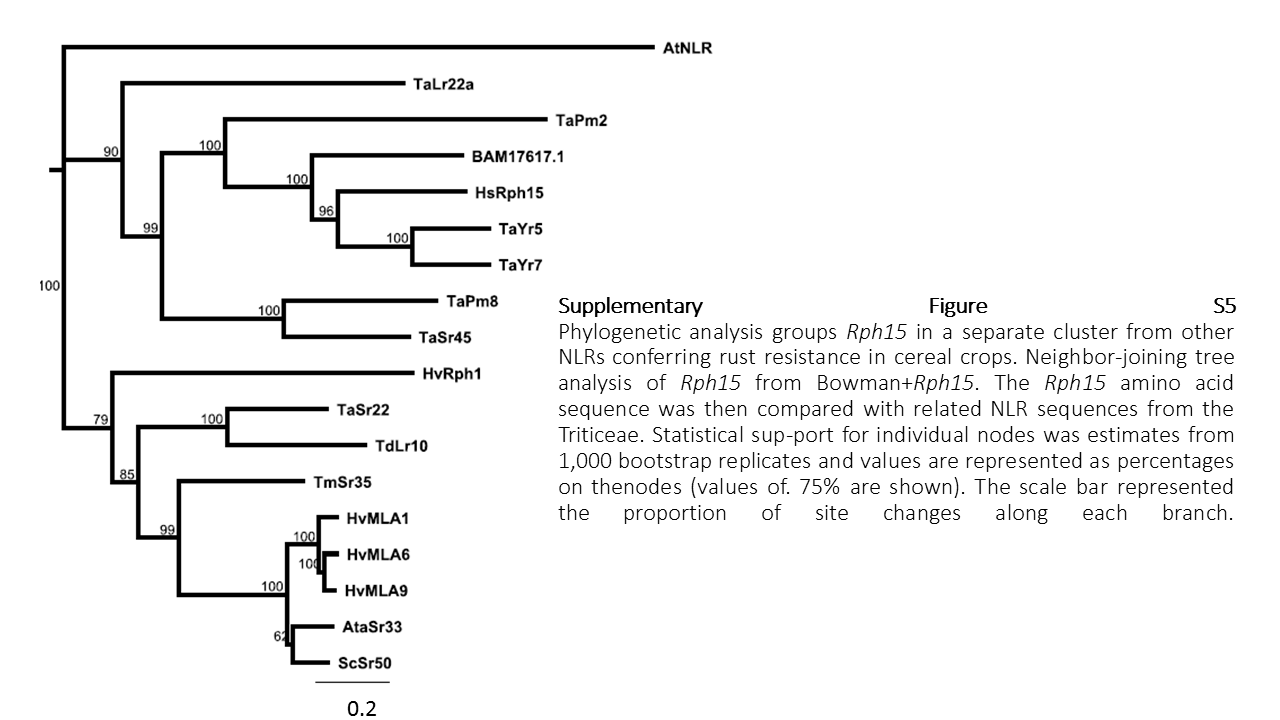
