## Supplementary Table 1 for "Ancient BED-domain-containing immune receptor from wild barley confers widely effective resistance to leaf rust"

**Supplementary Table 1- Phenotypic response testing at the seedling stage of barley leaf rust differential host genotypes using six pathogenically diverse North American *Puccinia hordei* isolates**

| **Differential host genotype** | **89-3** | **Neth28** | **90-3** | **92-7** | **92-6** | **90-5** |
| --- | --- | --- | --- | --- | --- | --- |
| Sudan (*Rph1*) | S | S | S | S | S | S |
| Peruvian (*Rph2*) | S | S | S | S | S | S |
| Estate (*Rph*3) | R | R | R | S | S | R |
| Gold (*Rph4*) | S | R | S | S | S | S |
| Magnificent (*Rph5*) | S | S | S | S | S | R |
| Bowman/Bol (*Rph6*) | S | S | S | S | S | S |
| Cebada capa (*Rph7*) | S | S | R | S | S | R |
| Egypt 4 (*Rph8*) | S | S | S | S | S | S |
| Hor2596 (*Rph9*) | S | R | S | S | R | S |
| Clipper BC8 (*Rph10*) | S | S | S | S | S | S |
| Clipper BC67 (*Rph11*) | S | S | S | S | S | S |
| Triumph (*Rph12*) | S | S | S | S | R | S |
| PI 531849 (*Rph13*) | R | R | R | R | R | S |
| PI 584760 (*Rph14*) | **R^a^** | **S^b^** | **S** | **S** | **R** | **S** |
| I95-282-2 (*Rph15*) | **R** | **R** | **S** | **R** | **R** | **R** |
| HS 680 (*Rph16*) | **R** | **R** | **S** | **R** | **R** | **R** |

^a^ Resistant and ^b^ Susceptible infection responses are highlighted yellow and red to denote isolate specificity between differential stocks carrying *Rph14*, *Rph15* and *Rph16* using the 6 North American *P. hordei* isolates.
