## Supplementary Table 2 for "Ancient BED-domain-containing immune receptor from wild barley confers widely effective resistance to leaf rust"

**Supplementary Table 2**- Genotypic results of 61 near isogenic lines carrying leaf rust resistance in the Bowman background published in Martin et al. (2020) with the gene-based *Rph15* KASP SNP marker developed in this study.

| **Line** | **Allele** | **Pedigree** | **KASP marker allele** |
| --- | --- | --- | --- |
| BW667 | *Rph15* | PI405338/3*Bowman | *Rph15* |
| BW668 | *?* | Bowman*4/PI405169A | allele other than Bowman and *Rph15* |
| BW669 | *Rph15* | PI405298/4*Bowman | *Rph15* |
| BW670 | *Rph15* | PI391007/4*Bowman | *Rph15* |
| BW671 | *Rph15* | PI466470/5*Bowman | *Rph15* |
| BW672 | *?* | Bowman*5//Moore/PI466373 | allele other than Bowman and *Rph15* |
| BW673 | *Rph15* | Bowman*6//Aim/PI405303 | *Rph15* |
| BW674 | *Rph15* | Bowman*7/PI466245 | *Rph15* |
| BW678 | *Rph15* | Bowman*4/PI354926 | *Rph15* |
| BW679 | *Rph15* | Bowman*6/PI282610 | *Rph15* |
| BW680 | *Rph15* | PI391049/4*Bowman | *Rph15* |
| BW681 | *Rph15* | PI391044/ND13944//4*Bowman | *Rph15* |
| BW688 | *Rph15* | PI354940/5*Bowman | *Rph15* |
| BW689 | *Rph15* | Bowman*5/PI405277 | *Rph15* |
| BW690 | *Rph15* | PI354923/5*Bowman | *Rph15* |
| BW691 | *Rph15+?* | Bowman*5/PI391045A het Eam1 | allele other than Bowman and *Rph15* |
| BW692 | *Rph15* | PI296841/6*Bowman | *Rph15* |
| BW693 | *Rph15* | PI354932/6*Bowman | *Rph15* |
| BW694 | *Rph15* | PI391002/6*Bowman | *Rph15* |
| BW695 | *Rph15* | PI391072/6*Bowman | *Rph15* |
| BW696 | *Rph15* | PI391087/6*Bowman | *Rph15* |
| BW697 | *Rph15+?* | Bowman*6/PI391121 | allele other than Bowman and *Rph15* |
| BW698 | *?* | Bowman*6/PI405154 | Bowman |
| BW699 | *Rph15* | PI405194/6*Bowman | *Rph15* |
| BW700 | *Rph15* | Bowman*6/PI405201 | *Rph15* |
| BW701 | *?* | PI405215/ND13944//5*Bowman | allele other than Bowman and *Rph15* |
| BW702 | *Rph15* | Bowman*6/PI405219 | *Rph15* |
| BW703 | *Rph15* | PI405233/6*Bowman | *Rph15* |
| BW704 | *Rph15* | PI405289/6*Bowman | *Rph15* |
| BW705 | *Rph15* | PI405308/6*Bowman | *Rph15* |
| BW706 | *Rph15* | Bowman*6/PI405332 | *Rph15* |
| BW707 | *Rph15* | Bowman*6/PI405335 | *Rph15* |
| BW708 | *Rph15* | Bowman*6/PI405364 | allele other than Bowman and *Rph15* |
| BW709 | *Rph15+?* | Bowman*6/PI405399 | *Rph15* |
| BW710 | *Rph15* | Bowman*7/PI405235 | *Rph15* |
| BW711 | *Rph15* | Bowman*7/PI405311 | *Rph15* |
| BW712 | *Rph15* | PI354928/4*Bowman | allele other than Bowman and *Rph15* |
| BW713 | *Rph15* | PI391000/6*Bowman | *Rph15* |
| BW714 | *Rph15* | PI391024/6*Bowman | *Rph15* |
| BW715 | *Rph15* | Bowman*6/PI355444 | *Rph15* |
| BW716 | *Rph15* | Bowman*6/PI405227 | allele other than Bowman and *Rph15* |
| BW717 | *Rph15* | PI405236/6*Bowman | *Rph15* |
| BW718 | *Rph15* | Bowman*6/PI405354A | *Rph15* |
| BW719* | *Rph15* | PI 355447/8*Bowman | *Rph15* |
| BW720 | *Rph15* | PI391089/8*Bowman | *Rph15* |
| BW721 | *Rph15* | Bowman*8/PI354937 | *Rph15* |
| BW722 | *Rph15* | Bowman*8/PI391069 | *Rph15* |
| BW723 | *Rph15* | PI405179/4*Bowman 0:cn | *Rph15* |
| BW724 | *Rph15* | Bowman*6/PI405341 | *Rph15* |
| BW725 | *Rph15* | PI391004/7*Bowman Translocation | *Rph15* |
| BW726 | *?* | HS 584/Bowman | *Rph15* |
| BW727 | *?* | HS 580/Bowman | Bowman |
| BW728 | *?* | Bowman*4/PI355445 | allele other than Bowman and *Rph15* |
| BW729 | *Rph15* | Bowman*5/PI355434 | *Rph15* |
| BW730 | *Rph15* | PI405210/5*Bowman | *Rph15* |
| BW733 | *Rph15* | PI405220/6*Bowman | allele other than Bowman and *Rph15* |
| BW734 | *Rph15* | PI466483/6*Bowman | *Rph15* |
| BW735 | *?* | 81882 *Rph17*.af/Bowman | Bowman |
| BW736 | *?* | Bowman*2/38P18-9-4 | Bowman |
| BW749 | *?* | Bowman*6/TUNISIA 33 | allele other than Bowman and *Rph15* |
| BW751 | *?* | 12022/3/CMB822/BOW//Columbia/4/6*Bowman | allele other than Bowman and *Rph15* |
| Controls |  | Bowman+*Rph15* (Park differential) | *Rph15* |
| Controls |  | Bowman+*Rph15* (Park differential) | *Rph15* |
| Controls |  | Bowman | Bowman |
| Controls |  | Bowman | Bowman |
| Controls |  | Bowman+*Rph15* (Park differential) plus Bowman | heterozygous call |
