## Supplementary Table 3 for "Ancient BED-domain-containing immune receptor from wild barley confers widely effective resistance to leaf rust"

| **Supplemenary Table 3- Summary table of rust pathogen isolates used in this study** | | | |
| --- | --- | --- | --- |
| **Pathogen** | **Disease** | **Pathotype name** | **avirulence/virulence spectra** |
| *Puccinia hordei* | Barley leaf rust | 5457 P+ | *Rph5,7,13,15,17,18,* 20, 21/1,2,3,4,6,8,9,12,19 |
|  |  | 90-3 | *Rph3,* 5, 7, 10, 11, 1, 17, 18/1, 2, 4, 6, 8, 9, 12, 14, 15, 16, 19 |
|  |  | 92-7 | *Rph11,* 13, 14, 15, 17, 18, 20/1, 2, 3, 4, 5, 6, 7, 8, 9, 10, 12, 16, 19 |
|  |  | 89-3 | *Rph3,* 13, 14, 15, 16/1, 2, 4, 5, 7, 8, 9, 10, 11, 12 |
|  |  | Neth28 | *Rph3,* 4, 9, 13, 15, 16/1, 2, 5, 6, 7, 8, 10, 11, 12, 14 |
|  |  | 92-6 | *Rph9,* 12, 13, 14, 15, 16/1, 2, 3, 4, 5, 6, 7, 8, 10, 11 |
|  |  | 90-5 | *Rph3,* 5, 7, 15, 16/1, 2, 4, 6, 8, 9, 10, 11, 12, 13 14 |
| *P. triticina* | Wheat leaf rust | 10-1,3,9,10,11,12 | *Lr3a,* 3bg, 3ka, 9, 11, 15, 17a, 18, 19, 21,23, 24, 25, 27+31, 28, 29, 30, 37/ Lr1, 2a, 10, 13, 14a, 16, 17b, 20, 26 |
| *P. graminis* f. sp. *tritici* | Wheat stem Rust | 98-1,2,3,5,6,7 | *Sr7b,* 8b, 9e, 13, 15, 21, 22, 24, 26, 27, 30, 31, 32, 33, 34, 35, 36, 38/ 5, 6, 8a, 9b, 11, 17 |
| *P. striiformis* f. sp. *tritici* | Wheat stripe rust | 134 E16 A+ | *Yr1,* 3, 4, 5, 10, 17, 33/ 2, 6, 7, 8, 9, 25, 73+74 |
