## Supplementary Table 4 for "Ancient BED-domain-containing immune receptor from wild barley confers widely effective resistance to leaf rust"

**Supplementary Table 4. Summary table of the primers used in this study**

| **Primer Name** | **Squence(5'→3')** | **Application** |
| --- | --- | --- |
| RGA4 F1 | CATTATCATAAGTGTAACCCCTGCAG | cloning of Rph15 |
| RGA4 R1 | TGCCGGATTATTTGCTCAGA | cloning of Rph15 |
| RGA4 F2 | CCCTTTGCTTGTGGGGACT | sequencing |
| RGA4 F3 | CATGGGATGATTTTGTCGACA | sequencing |
| RGA4 F4 | TGGGTGTCTAACAGCTTGGATG | sequencing |
| RGA4 F5 | TCTGGTCTAATGCATGATTTTGC | sequencing |
| RGA4 F6 | CTGAAAGTTCTAGCAGCGGTGT | sequencing |
| RGA4 F7 | AGAGCATGCCCTCCCATC | sequencing |
| RGA4 F8 new | ATTGAAATCCCTACAGCTGCACT | sequencing |
| RGA4 F9 | TCCACCATCAGTTTACGGTAAAAG | sequencing |
| RGA4 F10 | CGAAGAGATCAGAAGGTACGAGG | sequencing |
| XL-TOPO R | CACAGGAAACAGCTATGACCATG | sequencing |
| XL-TOPO F | AGGGTTTTCCCAGTCACGAC | sequencing |
| RGA4 F2 ATG | ATGGAGGACGCTTACCTTGTG | Mutant confirmation |
| Rph15 R-STOP | TCAGTGCACATATTGGTGGTCA | Mutant confirmation |
| Rph15 3'RACE -2 | TCAGGAAGTTGGGACTCTACAACAACCAGG | Rph15 RACE |
| Rph15 3'RACE-2 nest | TGTTACTGGAAGAATTGGATATTCGGGGC | Rph15 RACE |
| Rph15 5'RACE-2 | GTCCGCGAGAGATTTATCCAGCCGCGC | Rph15 RACE |
| Rph15 5'RACE-2 nest | ACCCCCTTTGCATCGGAAGCCAG | Rph15 RACE |
| Rph15 attB F new | GGGGACAAGTTTGTACAAAAAAGCAGGCTTAATGGAGGACGCTTACCTTGTG | cloning of Rph15-CC-BED-CC, Rph15-Full length, Rph15-CC-BED-CC-NB |
| Rph15 attBR1 new2 | GGGGACCACTTTGTACAAGAAAGCTGGGTCTCAGTGCACATATTGGTGGTCAATT | cloning of Rph15-NB-LRR, Rph15-Full length |
| Rph15 attb R2 new2 | GGGGACCACTTTGTACAAGAAAGCTGGGTCTCAGGCTGTACTTTGATGGTGACTTG | cloning of Rph15-CC-BED-CC |
| Rph15-Tr-attb-F1 | GGGGACAAGTTTGTACAAAAAAGCAGGCTTAatgtcagatccaagccgaagaa | cloning of Rph15-NB-LRR, Rph15-NB |
| Rph15-Tr-attb-R1 | GGGGACCACTTTGTACAAGAAAGCTGGGtcacaaatcagccagatagtccat | cloning of Rph15-NB and Rph15-CC-NB |
| Rph15-BD1-F | caagtttgtacaaaaaagcaggcttaatggatcatggcacattgaggggcgatcat | cloning of Rph15-BED-CC |
| Rph15-BD1-R | atgatcgcccctcaatgtgccatgatccattaagcctgcttttttgtacaaacttg | cloning of Rph15-BED-CC |
| Rph15-BD2-F | cagccgccaagccattcaagcatagatgcttgacccagctttcttgtacaaagtgg | cloning of Rph15-CC_BED and Rph15-BED |
| Rph15-BD2-R | ccactttgtacaagaaagctgggtcaagcatctatgcttgaatggcttggcggctg | cloning of Rph15-CC_BED and Rph15-BED |
